# NPF4.1 imports embryo-derived GA_4_ to the endosperm to promote seed germination

**DOI:** 10.1101/2024.10.07.617022

**Authors:** Mathilde Sirlin-Josserand, Lali Sakvarelidze-Achard, David Pflieger, Jean-Michel Davière, Patrick Achard

**Affiliations:** Institut de biologie moléculaire des plantes, CNRS, University of Strasbourg, 67084 Strasbourg, France

## Abstract

Gibberellins (GAs) are plant hormones essential for seed germination. In non-dormant seeds, humidity and light induce a fast increase in GA levels, which in turn enhances the growth potential of the embryo and weakening of the surrounding endosperm. While it is admitted that the tissue-specific distribution of GAs must be finely regulated in germinating seeds, it remains unclear where the bioactive forms are synthetized and how they are transported to their site of action. Here we demonstrate that bioactive GAs are mainly produced in the hypocotyl of the embryo and are transported in the endosperm, a few hours from the onset of imbibition, to induce the expression of cell wall loosening genes such as *EXPANSIN2*. We further show that NITRATE TRANSPORTER1/PEPTIDE TRANSPORTER FAMILY (NPF) NPF4.1, previously identified as a cellular GA influx transporter, is localized in the plasma membrane of endosperm cells and contributes in the transport of embryo-derived GA_4_ into the endosperm. Accordingly, *npf4.1* mutant seeds are hypersensitive to the GA biosynthesis inhibitor paclobutrazol in germination assay, and the phenotype is suppressed by application of exogenous bioactive GAs. Finally, we show that two other GA transporters, NPF3.1 and NPF2.13, are expressed in the embryo of germinating seeds and may modulate cellular GA levels in elongating cells of the radicle and cotyledons.

## Introduction

Seed germination is a critical phase in the plant life cycle and the first developmental process leading to plant establishment. This phase is tightly controlled by a set of environmental cues, mainly moisture, light and temperature, which are integrated and transmitted by internal hormonal signals (1). Among phytohormones, gibberellins (GAs) are tetracyclic diterpenoid molecules known for long time to promote seed germination (2). This is exemplified by the defect of GA-deficient mutant seeds that fail to germinate without exogenous GA treatment (3). A similar alteration can be observed by applying inhibitors of GA biosynthesis, demonstrating that *de novo* GA synthesis is required for the accomplishment of seed germination (4). GAs promote seed germination by stimulating the degradation of repressing DELLA proteins (DELLAs), through the ubiquitin-proteasome pathway (5–7). DELLAs belong to a small group of nuclear regulators that interact with and modulate the activity of transcription factors, thereby controlling gene expression (7). The *Arabidopsis* genome contains five *DELLA* genes and previous studies have reported that GAs regulate seed germination via suppression of the DELLA RGA-LIKE2 (RGL2) function (6, 8–10). Genetic analyses showed that the absence of RGL2 integrally restores seed germination in GA-deficient mutants (6, 8).

GA/RGL2 signaling regulates seed germination by at least two complementary mechanisms. First, it increases the growth potential of the embryo (radicle–hypocotyl growth zone), triggering mechanical forces that contribute to the rupture of the surrounding coats, which include living endosperm and dead testa (11, 12). This is principally achieved by cell expansion along the embryo axis. Immediately after imbibition, GA signaling induces in the radicle the expression of cell wall remodeling enzyme (CWRE) genes, such as *EXPANSIN8* (*EXP8*), first in the epidermis and then in the inner layers to coordinate radicle tip growth (13). Later on, GA signaling induces cell expansion in the hypocotyl, increasing the overall growth potential of the embryo (14). Secondly, GAs also play an essential role in weakening the seed coats surrounding the embryo, which provides a physical constraint to the protrusion of the radicle (12, 15, 16). Previous work has shown that GA signaling increases the activity of two NAC transcription factors (NAC25 and NAC1L) that induce in endosperm the expression of *CWRE* genes such as *EXPANSIN2* (*EXPA2*) (17). In doing so, GAs positively regulate the expansion and separation of endosperm cells to allow embryo growth and radicle protrusion. Taken together, these findings demonstrate that GA levels must be tightly regulated in space and time during seed germination.

The spatio-temporal distribution of biologically active GAs (mostly GA_4_ in *Arabidopsis*; 18) is regulated at several levels including hormone metabolism and transport (19, 20). Previous studies have reported that while GA levels are low in dry mature seeds, the levels of GA_4_ increase rapidly after imbibition and exposure to light (21, 22). In agreement with this finding, the expression of GA 20-oxidase (GA20ox) and GA 3-oxidase (GA3ox) gene families, involved in the latest GA biosynthetic steps (Figure 1a), increases substantially after seed imbibition (22). The fact that *GA3ox1* and *GA3ox2* transcripts accumulate predominantly in the embryo, it was then proposed that bioactive GA_4_ moves from the embryo to the endosperm (21). Interestingly, a similar finding was made in germinating barley grains, where bioactive GAs produced in the embryo move towards the aleurone layers of the endosperm to induce the expression of α-amylase required for starch hydrolysis (23, 24). This spatial separation between the production and needed site emphasizes the existence of an active transport of GAs within the seed. In recent years, several plasma membrane-localized GA transporters have been characterized and shown to concentrate GAs in specific cell types or tissues in the plant (7, 19). For example, the NITRATE TRANSPORTER1/PEPTIDE TRANSPORTER family (NPF) gene *NPF3.1* encodes a GA transporter that concentrates bioactive GAs in elongating root endodermis, the primary GA-responsive tissue controlling root growth (25, 26). Since then, several GA transporters have been identified through *in vitro* screening in yeast and *Xenopus* oocyte (based on cellular GA uptake), but only a few have been confirmed *in planta* (27, 28). Interestingly, three GA transporters have been reported to play a role during seed germination: NPF3.1, SWEET13 and SWEET14 (29, 30). However, despite the fact that *npf3.1* and *sweet13 sweet14* loss-of-function mutants display altered responses to GA during germination, it is unknown where these GA transporters act in germinating seeds. Thus at present, if it is accepted that GAs are essential for the completion of seed germination, it remains unclear where the bioactive GAs are produced, and how and when they are distributed in needed tissues. Here, we report that bioactive GAs are essentially produced in the embryo and are required in the endosperm to promote its rupture. In non-dormant seeds, we demonstrate that bioactive GA_4_, produced in the embryo, begins to move towards the endosperm between 6 to 9 hours after the onset of imbibition. Accordingly, we show that embryo-derived GA_4_ induces the expression of *EXPA2* in endosperm. Furthermore, we demonstrate that NPF4.1, localized at the plasma membrane of endosperm cells, facilitates the transport of GA_4_ from the embryo to the endosperm. Finally, we discuss the role of NPF3.1 and NPF2.13 during seed germination.

**Figure 1.**
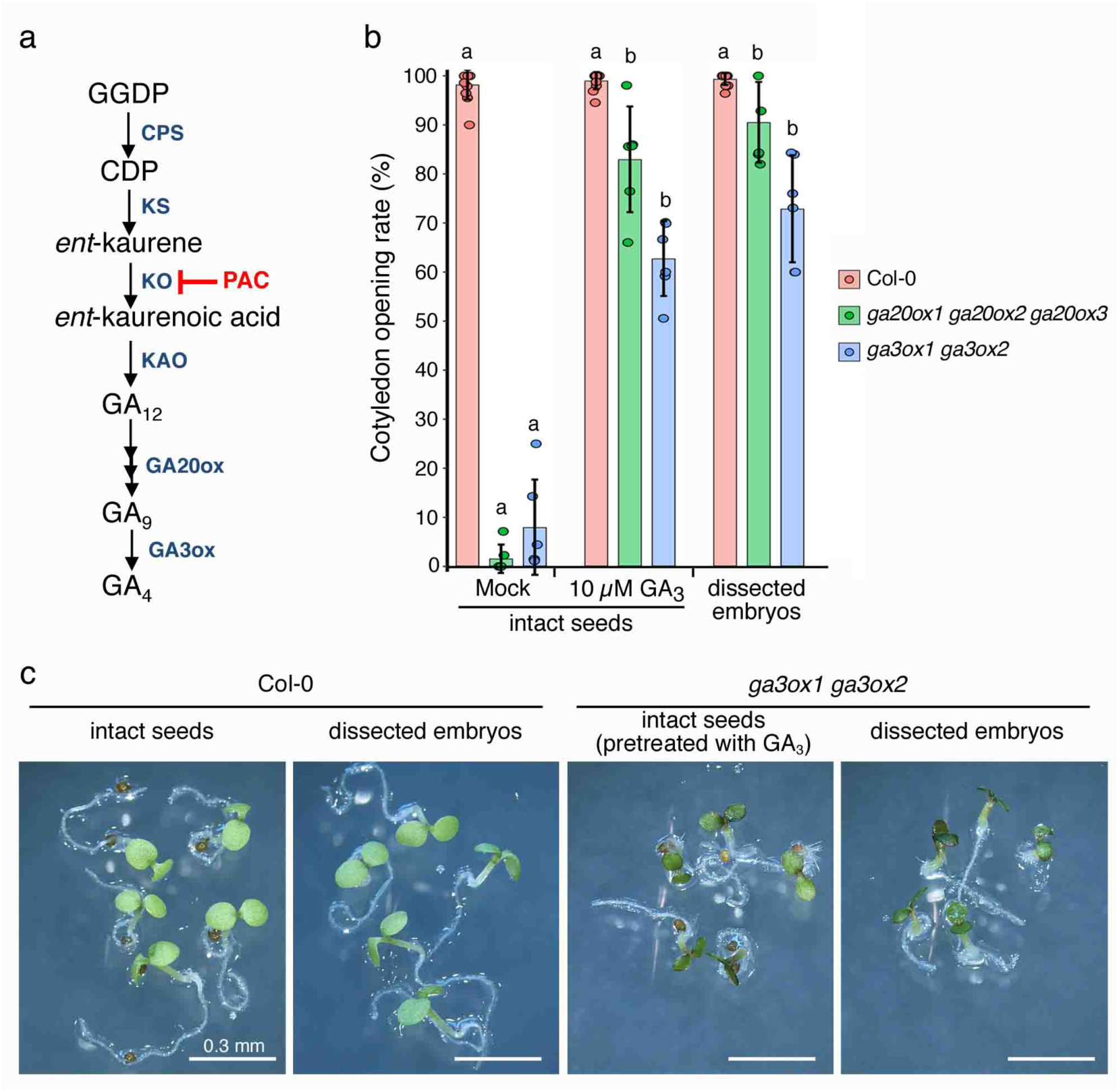
GA-induced weakening of seed coats is a limiting step in germination process. (a) GA biosynthetic pathway. GA biosynthetic enzymes are indicated in blue. Paclobutrazol (PAC), chemical inhibitor of *ent-*kauren oxidase (KO) activity. GGDP, geranyl geranyl diphosphate; CDP, *ent*-copalyl diphosphate; KS, *ent*-kauren synthase; KAO, *ent*-kaurenoic acid oxidase; GA20ox, GA 20-oxidase; GA3ox, GA 3-oxidase. (b) Germination rate of wild-type (Col-0), *ga20ox1 ga20ox2 ga20ox3* and *ga3ox1 ga3ox2* mutant seeds treated with 10 µM GA_3_ and mock controls, or dissected embryos separated from their seed coats. Germination was determined as cotyledon opening after 4 days of imbibition. The values are means ±SD of two independent experiments using three seed batches of approximately 50 seeds. Different letters denote significant differences (p<0.05) using one-way ANOVA with Tukey’s test for multiple comparisons. (c) Representative phenotypes of 4-day-old wild-type and *ga3ox1 ga3ox2* seedlings originating from intact seeds or dissected embryos. *ga3ox1 ga3ox2* intact seeds were pretreated with GA_3_ during the stratification. Scale bar, 0.3 mm.

## Results

### GA-induced weakening of seed coats is a limiting step for seed germination

Seed germination is a multi-step process that begins with the absorption of water and ends with the emergence of the radicle through the seed coats (1). Given that GAs fulfill multiple roles during germination, we carried out germination tests with GA biosynthesis mutants, in order to gain a better understanding of the territories where GA activity is mostly required. To this end, we compared the germination percentage of intact seeds and dissected embryos (the surrounding envelopes are removed mechanically) of wild-type (Col-0), *ga20ox1 ga20ox2 ga20ox3* and *ga3ox1 ga3ox2* mutants. The triple *ga20ox1-2-3* and double *ga3ox1-2* mutant seeds are unable to produce bioactive GAs and as a consequence, fail to germinate without exogenous GAs (Figure 1a,b; 31, 32). In this assay, to be able to compare the germination rate of intact and dissected seeds, we did not measure the completion of the germination (protrusion of the radicle), but the opening of the cotyledons, 4 days after imbibition. Remarkably, we found that GA treatment or removal of the seed coats substantially restored the germination defect of triple *ga20ox1-2-3* and double *ga3ox1-2* mutant seeds, as previously reported with the GA-deficient *ga1-3* mutant (Figure 1b; 33). Moreover, despite the absence of bioactive GAs, dissected *ga3ox1 ga3ox2* embryos gave rise to dwarf seedlings with open and green cotyledons, similar to those of intact GA-deficient seeds pretreated with GA (Figure 1c). Taken together, these results suggest that, although GAs are important for embryo growth, GA-induced weakening of the seed coats is the limiting step in the germination process.

### Bioactive GAs are synthetized in embryo during germination

In a subsequent experiment, we investigated where the bioactive GAs are produced in germinating seeds. Since GA3ox catalyzes the final step in the GA biosynthetic pathway (Figure 1a), the expression pattern of the corresponding genes provides valuable information on the main sites of bioactive GA production in the seed. Previous studies showed that *GA3ox1* and *GA3ox2* are expressed to high levels in the cortex and endodermis of hypocotyl embryos isolated from 16 to 24 hours imbibed seeds (21, 31). Thus, if it becomes clear that bioactive GAs are abundantly produced in the embryo, we have little information on their synthesis in the endosperm. To obtain an overview of the GA biosynthesis sites in germinating seeds, we took advantage of a recent transcriptomic analysis performed on embryos and endosperms isolated from seeds imbibed for 24 hours at 30°C (34). Comparison analysis revealed that while the early steps of GA biosynthesis occur equally in both seed compartments, *GA3ox* transcripts are substantially more abundant in the embryo than in the endosperm, confirming that bioactive GAs are mainly produced in the embryo (Supplementary Figure 1). Moreover, it also confirmed that *GA3ox1* and *GA3ox2* are the two main *GA3ox* genes expressed in germinating seeds (only few *GA3ox4* transcripts were detected), and it explains why the *ga3ox1 ga3ox2* double mutant fails to germinate. Noteworthy, we also found that *GA 2-oxidase* (*GA2ox*) catabolism genes are globally more expressed in the endosperm than in the embryo, underlying a rapid GA turnover in the endosperm (Supplementary Figure 1).

We then investigated the temporal expression patterns of selected *GA20ox* and *GA3ox* genes along the germination process. In our experimental conditions (see Methods), non-dormant Col-0 seeds show rupture of the seed coats between 12 and 24 hours after imbibition, and the protrusion of the radicle between 24 and 36 hours after imbibition (Figure 2a,b). With the aim of analyzing GA biosynthetic gene expression levels ahead of each of these two developmental stages, total RNA was extracted from endosperms and embryos dissected 9 and 18 hours after seed imbibition. As controls, we used two gene markers, *ABA INSENSITIVE4* (*ABI4*) and *EXPA2* that are strictly expressed in embryo and endosperm, respectively (Figure 2c; 35, 36). In line with the above results, *GA20ox3*, *GA3ox1* and *GA3ox2* are the three most expressed GA biosynthetic genes in germinating seeds (Figure 2c). *GA20ox3* is mostly expressed in the endosperm, while *GA3ox1* and *GA3ox2* are more highly expressed in the embryo, at both 9 and 18 hours after imbibition. By contrast, *GA20ox1* and *GA20ox2* are expressed at low and homogenous levels in embryo and endosperm (Figure 2c). Therefore, if it is likely that inactive GA_9_ (product of the GA20ox) accumulates in both seed compartments before the rupture of the endosperm and the protrusion of the radicle, the bioactive GA_4_ is mainly produced in the embryo.

**Figure 2.**
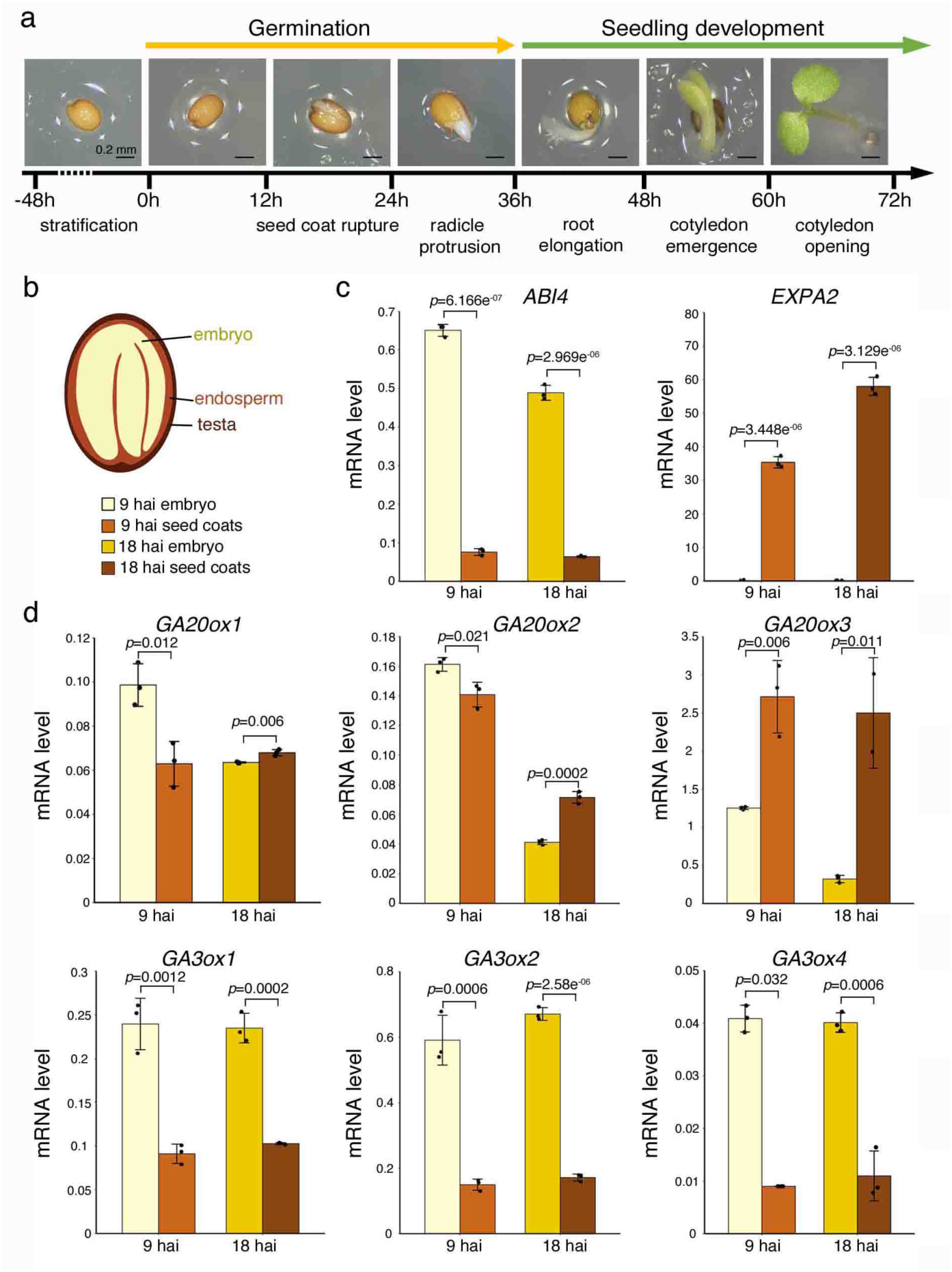
Temporal and spatial expression of GA biosynthetic genes in germinating seeds. (a) Morphology of *Arabidopsis* seeds at several different developmental stages after imbibition. After stratification of the seeds (breaking of dormancy), germination begins with water uptake and is completed with the protrusion of the radicle, which is followed by the postgerminative growth of the seedling. Approximate time of each developmental stage in our growing conditions is indicated. Scale bar, 0.2 mm. (b) Schematic representation of a mature *Arabidopsis* seed, consisting of an embryo surrounded by seed coats (a living endosperm and dead testa). hai, hours after imbibition starting point. (c) Expression level of *ABI4* (embryo marker), *EXPA2* (endosperm marker) and selected *GA20ox* and *GA3ox* gene transcripts in separated embryos (yellow) and seed coats (brown) isolated from 9 hours and 18 hours imbibed seeds (hai). Data are means ±SD of three technical replicates. *P*-values are from Student’s t test. Similar results were obtained in two independent experiments.

To further substantiate the tissue-specific expression patterns of GA biosynthetic genes in germinating seeds, we used *GA20ox*- and *GA3ox-GUS* reporter lines. GUS staining was performed on dissected embryos and seed coats, 18 hours after seed imbibition, so before radicle protrusion. Overall, the results corroborate with the above gene expression analyses; GA3ox1-GUS and GA3ox2-GUS are only detected in the embryo hypocotyl and GA20ox3-GUS is detected in both embryo and endosperm (Figure 3). Also, GA20ox1-GUS and GA20ox2-GUS are detected in the radicle while *GA3ox4-GUS* line shows no GUS activity in the seed. Altogether, these different approaches strongly suggest that *GA20ox* and *GA3ox* are expressed in germinating embryos, supporting the role of GAs in the increased growth potential of the embryonic axis. On the contrary, *GA3ox* expression is low, if not excluded from the endosperm.

**Figure 3.**
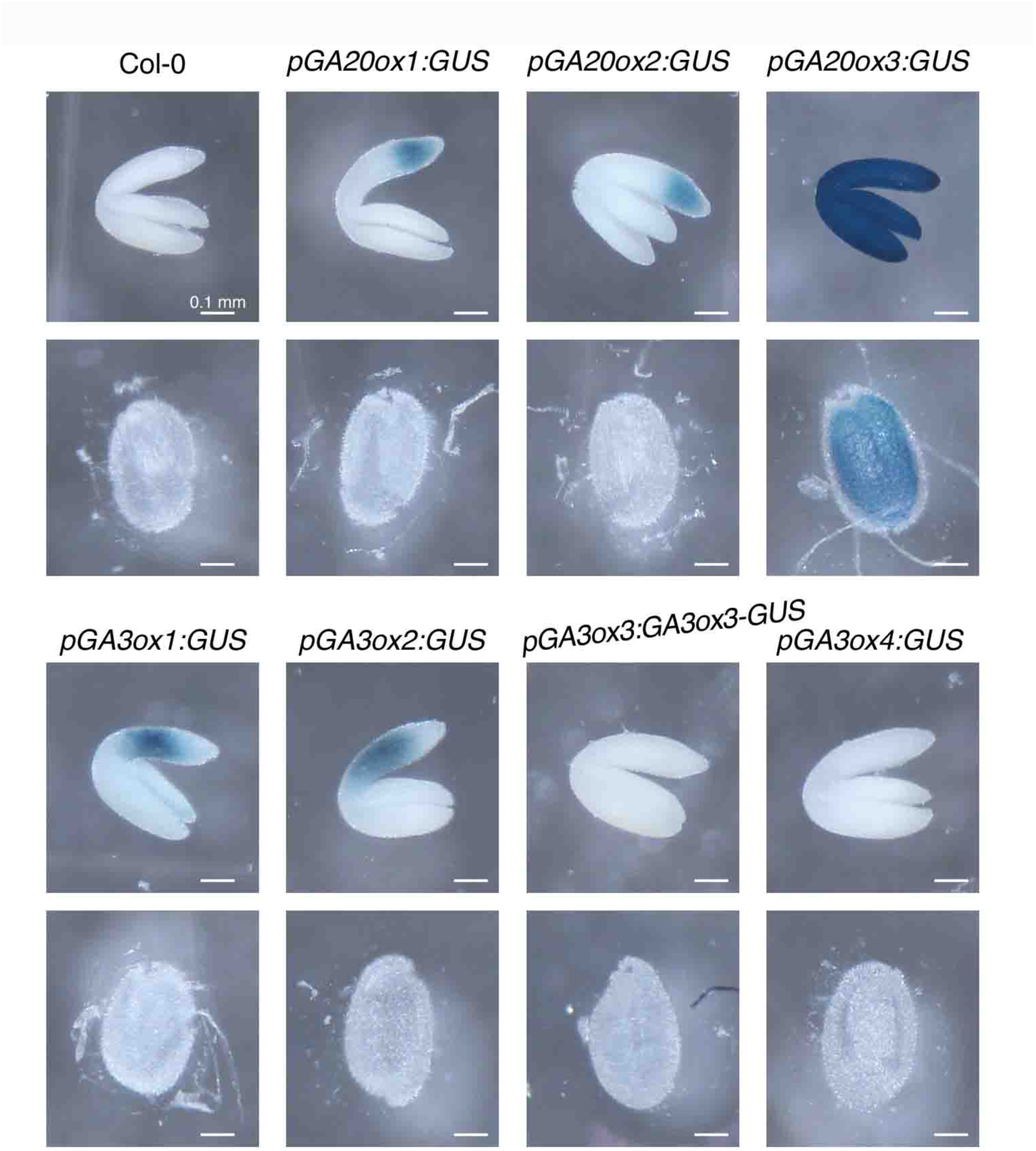
Tissue-expression of *GA20ox* and *GA3ox* genes. Comparison analysis of *GA20ox:GUS* and *GA3ox:GUS* expression patterns in separated embryos (top panels) and seed coats (bottom panels) isolated from 18 hours imbibed seeds. Similar results were obtained in two independent experiments. Scale bar, 0.1 mm.

### GA_4_ is transported from embryo to endosperm

We have previously shown that GA-induced weakening of seed coats is a limiting step for germination, and that *GA3ox* expression is mostly restricted to the embryo. It is therefore conceivable that bioactive GAs or a GA-induced mobile signal produced by the embryo, are transmitted in the endosperm. To study in more detail the nature of this embryo-to-endosperm signal, we monitored in different contexts and time prior to endosperm cell separation, the spatial expression of *EXPA2* (by promoter GUS studies), an endosperm-specific CWRE marker of cell expansion that is induced by GAs (17). Consistently, as a control, we also showed that *EXPA2* expression is restricted to endosperm and depends on an embryo signal that is emitted between 4 and 24 hours after imbibition, in our experimental conditions (Supplementary Figure 2a). In addition, we also confirmed that GAs induce the expression of *EXPA2* (Supplementary Figure 2b).

To assess the nature of the embryo-derived signal inducing *EXPA2* expression, we drew inspiration from the “seed coat bedding” assay developed by Lee et al. (2010)(37), which consists of assembling dissected embryos and seed coats from different genotypes. In our assay, we placed separated *pEXPA2:GUS* seed coats on a layer of Col-0 or *ga3ox1 ga3ox2* embryos, dissected from seeds imbibed for 1 hour (Figure 4a). 24 hours after incubation, seed coats were assayed for GUS activity. Remarkably, we detected a strong EXPA2:GUS activity only in the endosperm of seed coats incubated with Col-0 embryos (Figure 4b). This result confirms that the signal released by the embryo is a GA-regulated molecule and strongly suggests that it is a bioactive GA.

**Figure 4.**
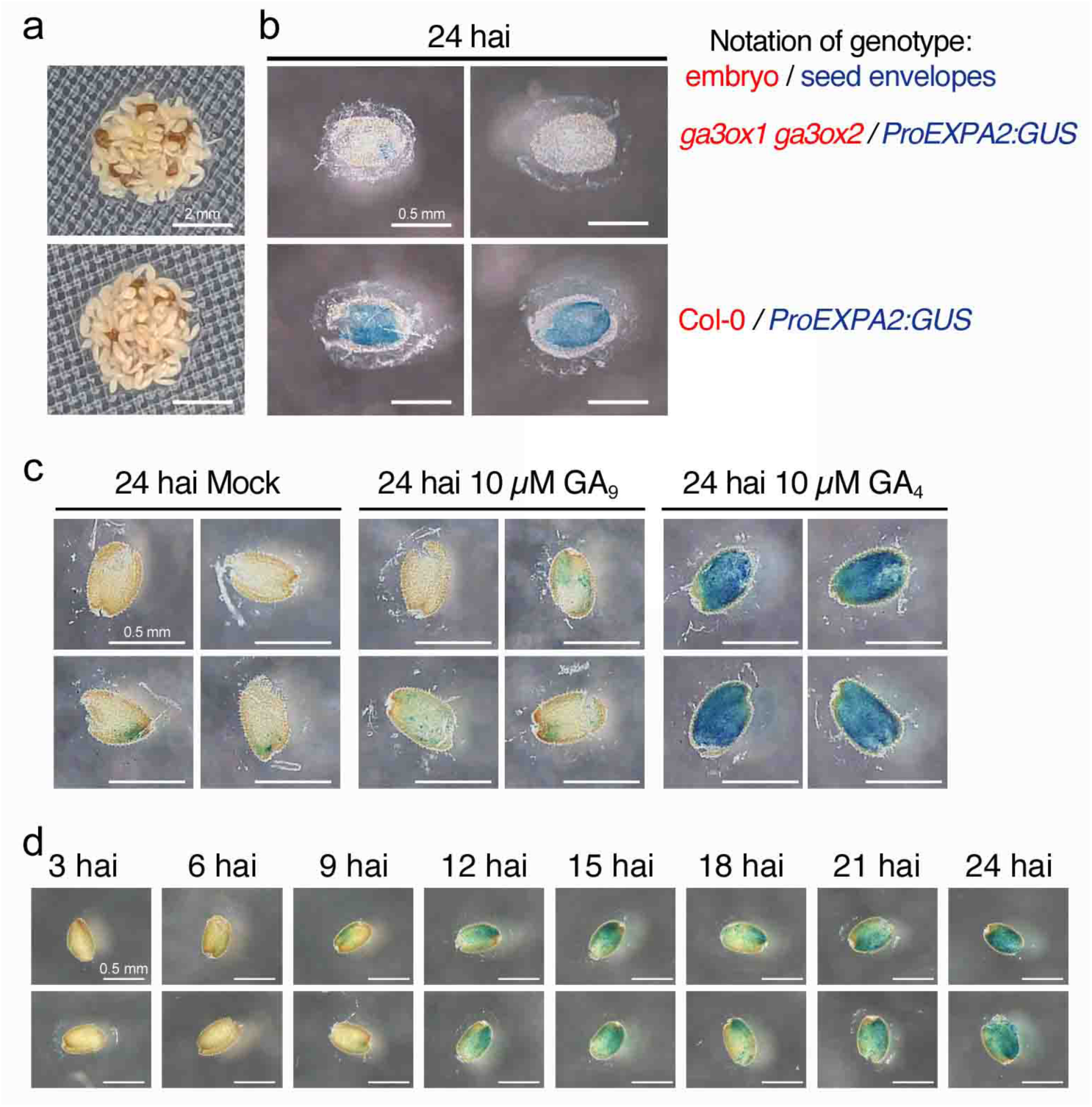
GA_4_ is transported from embryo to endosperm shortly after imbibition. (a) Embryo bedding assays using separated *pEXPA2:GUS* seed coats disposed on a layer of separated wild-type or *ga3ox1 ga3ox2* embryos, isolated one hour after imbibition. Scale bar, 2 mm. (b) Pictures show EXPA2:GUS staining in endosperm, performed 24 hours after incubation of the seed coats with the embryos. Scale bar, 0.5 mm. (c) EXPA2:GUS staining of dissected seed coats, isolated one hour after seed imbibition, incubated with 10 µM GA_9_ or 10 µM GA_4_ (and controls, Mock) for 24 hours. Similar results were obtained in two independent experiments. Scale bar, 0.5 mm. (d) Induction kinetics of EXPA2:GUS activity in imbibed seed coats. Seed coats from *pEXPA2:GUS* seeds imbibed from 3 to 24 hours were separated. EXPA2:GUS activity was then assayed on the separated seed coats. Similar results were obtained in two independent experiments. Scale bar, 0.5 mm. hai, hours after imbibition.

To strengthen this result, a complementary analysis was carried out using exogenous GAs. Seed coats separated from *pEXPA2:GUS* seeds previously imbibed for 1 hour, were incubated with 10 µM GA_9_ (substrate of GA3ox) or 10 µM GA_4_ (active form), or Mock as a control, for 24 hours. The results clearly show that solely GA_4_ is able to induce significant EXPA2:GUS activity in endosperm (Figure 4c). Taken together, these results confirm that GA_4_ represents the signaling molecule produced by the embryo, which is transported into the endosperm and induces the expression of *EXPA2*, necessary for endosperm rupture.

Finally, we defined the time interval after imbibition from which the transport of GA_4_ from embryo to endosperm is active. To this end, seed coats from *pEXPA2:GUS* seeds were separated from embryos at different time points after seed imbibition (from 3 to 24 hours) and subsequently incubated for a total period of 24 hours before that GUS activity was assayed. We started to detect weak EXPA2:GUS signal after 9 hours of imbibition (around the micropylar region) and then a more intense staining from 12 hours after imbibition (Figure 4d). These results thus indicate that GA_4_ produced in the embryo begins to be transported in the endosperm shortly before 9 hours of imbibition, prior to the induction of *EXPA2:GUS* in endosperm.

### Characterization of GA transporters acting in germinating seeds

To identify the transporters involved in the translocation of GA_4_ from embryo to endosperm, we examined in germinating seeds the expression levels of all known transporters displaying GA transport activity (27, 28, 30, 38). As above with the GA biosynthetic genes, we initially relied on the RNAseq analysis performed on separated embryos and endosperms, isolated from 24 hours imbibed seeds (34). Comparison analysis revealed that three transporters are substantially expressed in germinating seeds: *NPF2.13*, *NPF3.1* and *NPF4.1* (Supplementary Figure 3). Of note, these three transporters are highly expressed in the first few hours after imbibition, thus prior endosperm rupture (Supplementary Figure 4). Interestingly, *NPF4.1*, which encodes an active GA importer (25,39), is preferentially expressed in the endosperm and is therefore a strong candidate for the transport of embryo-derived GA_4_ into the endosperm (Supplementary Figure 3). To further substantiate these results, we determined the expression levels of *NPF2.13*, *NPF3.1* and *NPF4.1* in isolated embryos and endosperms, at the time when GA_4_ begins to be transported into the endosperm (e.g. 9 hours after imbibition). Consistent with the transcriptomic data, we found that *NPF3.1* is more expressed in embryos than in endosperms, and reciprocally, *NPF2.13* and especially *NPF4.1* are more expressed in the endosperm (Figure 5a). To complete the analysis, we also monitored their tissue-expression pattern in transgenic promoter:GUS transcriptional fusion lines. 9 hours after seed imbibition, NPF3.1:GUS activity was restricted to the radicle, NPF4.1:GUS activity was detected in the whole endosperm and NPF2.13:GUS activity was visible only in a few cells in cotyledons (boundary zone) and endosperm (Figure 5b).

**Figure 5.**
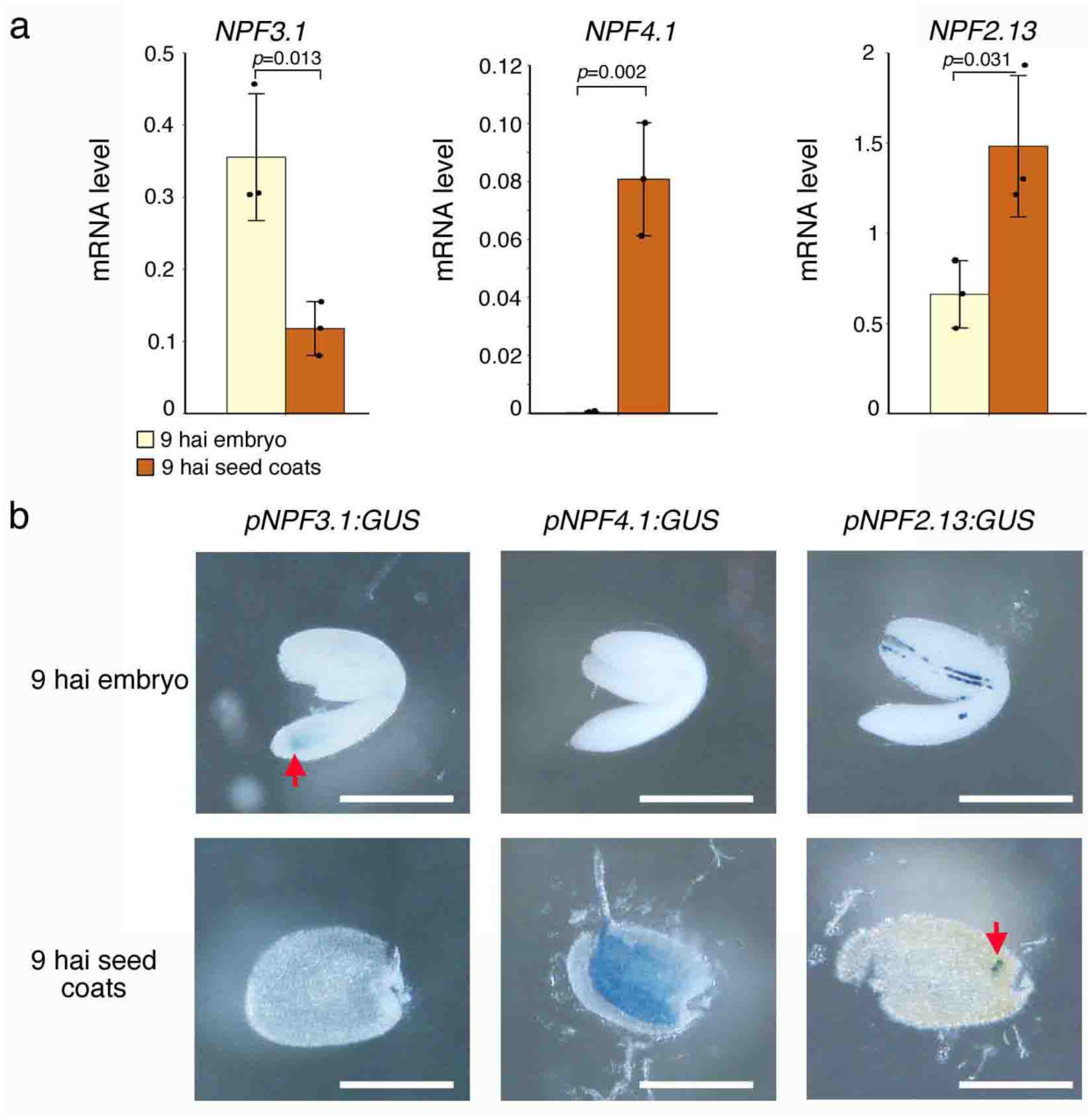
Spatial expression patterns of *NPF3.1*, *NPF4.1* and *NPF2.13* in imbibed seeds. (a) Expression level of *NPF3.1*, *NPF4.1* and *NPF2.13* in separated embryos (yellow) and seed coats (brown) isolated from 9 hours imbibed seeds. Data are means ±SD of three technical replicates. *P*-values are from Student’s t test. Similar results were obtained in two independent experiments. hai, hours after imbibition. (b) Expression patterns of *NPF3.1*, *NPF4.1* and *NPF2.13* in separated embryos and seed coats isolated from 9 hours imbibed *pNPF3.1:GUS*, *pNPF4.1:GUS* and *pNPF2.13:GUS* seeds. Similar results were obtained in two independent experiments. Red arrows indicate stained cells. Scale bars, 0.5 mm.

Finally, we addressed the subcellular localization of NPF4.1; NPF2.13 and NPF3.1 were shown to localize in the plasma membrane (25, 38). To this end, *p35S:NPF4.1-GFP* transgenic lines were generated and NPF4.1-GFP fluorescence was analyzed in root cells by confocal microscopy (Supplementary Figure 5). Similar to NPF2.13 and NPF3.1, NPF4.1 localized to the plasma membrane, which is consistent with previous work showing that *NFP4.1* expressing oocytes are able to import GAs (25, 39). Thus NPF4.1 is a GA importer involved in intercellular GA transport.

Taken together, these results indicate that NPF4.1 is a GA transporter that accumulates in the plasma membrane of endosperm cells, a few hours after seed imbibition, and could therefore contribute to the transport of bioactive GA_4_ from the embryo to the endosperm. By contrast, based on their tissue expression patterns, NPF2.13 and NPF3.1 are likely involved in the cellular distribution of GAs in the embryo.

### NPF4.1 facilitates embryo-to-endosperm GA transport, promoting seed germination

To assess the functional role of NPF4.1 during seed germination, we characterized two independent insertion lines, a loss of function mutant (*npf4.1-1*) and a knock down mutant (*npf4.1-2*), and generated two independent transgenic overexpression lines (*p35S:NPF4.1-HA* L4 and L8) (Supplementary Figure 6). In germination assays, non-dormant wild-type and *npf4.1* mutant seeds tended to germinate similarly under control conditions, all reaching 90 to 100% of germination two days after imbibition (Figure 6a). Interestingly, in the presence of low concentration of paclobutrazol (2 µM PAC), a GA biosynthesis inhibitor (40; Figure 1a), the germination rate of *npf4.1* mutant seeds was substantially lower than that of wild-type. 5 days after imbibition, the germinate rate of *npf4.1-1* and *npf4.1-2* mutant seeds was only of 34% and 20%, respectively, compared to 100% for the wild-type (Figure 6a). This hypersensitivity to PAC indicates that *npf4.1* mutants have reduced capacity to import endogenous GAs in endosperm. Accordingly, application of bioactive GAs significantly attenuated the effect of PAC for the *npf4.1* mutants (Figure 6a). Remarkably, as previously reported with other GA transporters (25), overexpression of *NPF4.1* also restrained the germination rate (Figure 6b). The effect was exacerbated in the presence of PAC, and application of GA only slightly increased the germination rate (Figure 6b). As previously suggested, it is likely that overexpression of *NPF4.1* affects its tissue distribution and causes retention of GAs at sites of synthesis, thus preventing movement of bioactive GAs from embryo to endosperm.

**Figure 6.**
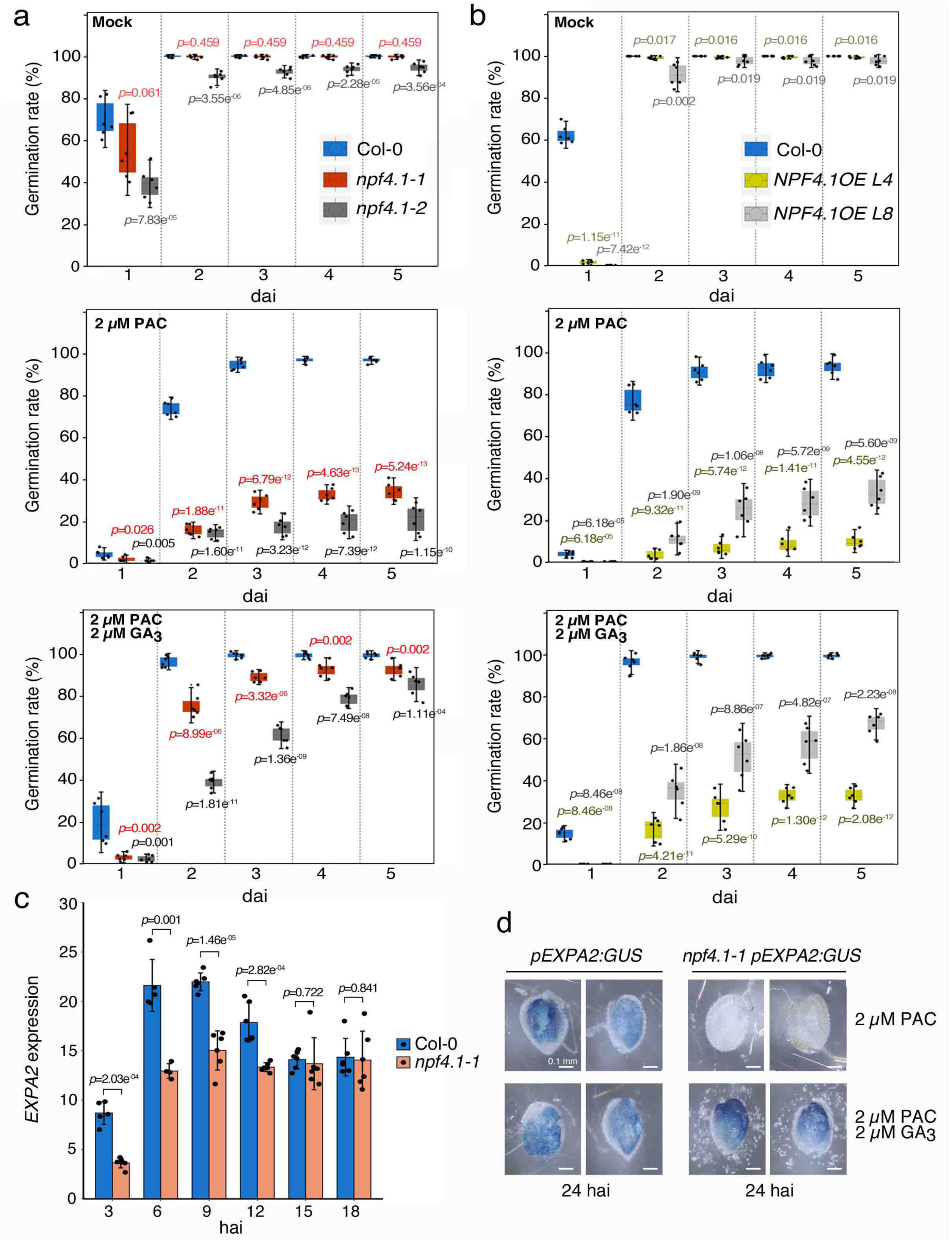
NPF4.1 activity modulates GA sensitivity during germination. (a,b) Germination of wild-type (Col-0), *npf4.1-1*, *npf4.1-2* and *p35S:NPF4.1-HA* (*NPF4.1OE* L4 and L8) seeds in the absence (Mock) or presence of 2 µM PAC or 2 µM PAC with 2 µM GA_3_, as indicated. Germination was determined as radicle protrusion from day 1 to day 5 after imbibition. The values are means ±SD of three seed batches of 50 to 100 seeds sown in duplicate. *P*-values are from Student’s t test for *npf4.1* or *NPF4.1OE versus* Col-0. dai, days after imbibition. (c) *EXPA2* expression level in imbibed wild-type (Col-0) and *npf4.1-1* mutant seeds imbibed for 3 to 18 hours. Data are means ±SD of two independent replicates with two to three technical replicates. *P*-values are from Student’s t test. hai, hours after imbibition. (d) EXPA2:GUS activity in dissected seed coats of *pEXPA2:GUS* and *npf4.1-1 pEXPA2:GUS* seeds, isolated one hour after seed imbibition, and incubated for 24 hours with 2 µM PAC or 2 µM PAC with 2 µM GA_3_. Similar results were obtained in two independent experiments. hai, hours after imbibition. Scale bars, 0.1 mm.

To evaluate more directly the role of NPF4.1 on the transport of GAs from the embryo to the endosperm, we analyzed the induction kinetics of *EXPA2* in wild-type and *npf4.1-1* imbibed seeds. Although the expression profile was similar, the transcript abundance of *EXPA2* was significantly lower in *npf4.1-1* mutant compared with the wild-type, especially from 3 to 12 hours after imbibition (Figure 6c). To further substantiate this result, we introgressed the *pEXPA2:GUS* construct into the *npf4.1-1* mutant. In contrast to wild-type seeds, no GUS activity was detected in *npf4.1* mutant background seeds imbibed with PAC for 24 hours. Moreover, GA application could entirely suppress the effect of PAC on EXPA2:GUS activity in *npf4.1-1* (Figure 6d). Thus, in agreement with the germination defect of *npf4.1* mutants on PAC (Figure 6d), these results demonstrate that NPF4.1 facilitates the translocation of embryo-derived GA_4_ into the endosperm, an important step in the regulation of germination.

### NPF2.13 and NPF3.1 regulate seed germination

We also investigated the biological function of NPF2.13 and NPF3.1 in germination assays. Similar to *npf4.1* mutants, the germination rate of *npf2.13* and *npf3.1* mutant seeds was significantly reduced in the presence of PAC compared to wild-type (Figure 7a,b), a response already observed with the *npf3.1* mutant (29). However, the sensitivity to PAC was less important in these two mutants than in the *npf4.1-1* mutant (48% and 54% of germination for *npf3.1* and *npf2.13*, respectively, and 15% for *npf4.1-1*, two days after imbibition; Figures 6b and 7b,e). As expected, application of bioactive GAs largely suppressed the effect of PAC (Figure 7a,b). This last result indicates that the phenotype of *npf2.13* and *npf3.1* mutants is associated with a defect in GA transport in the germinating seed.

**Figure 7.**
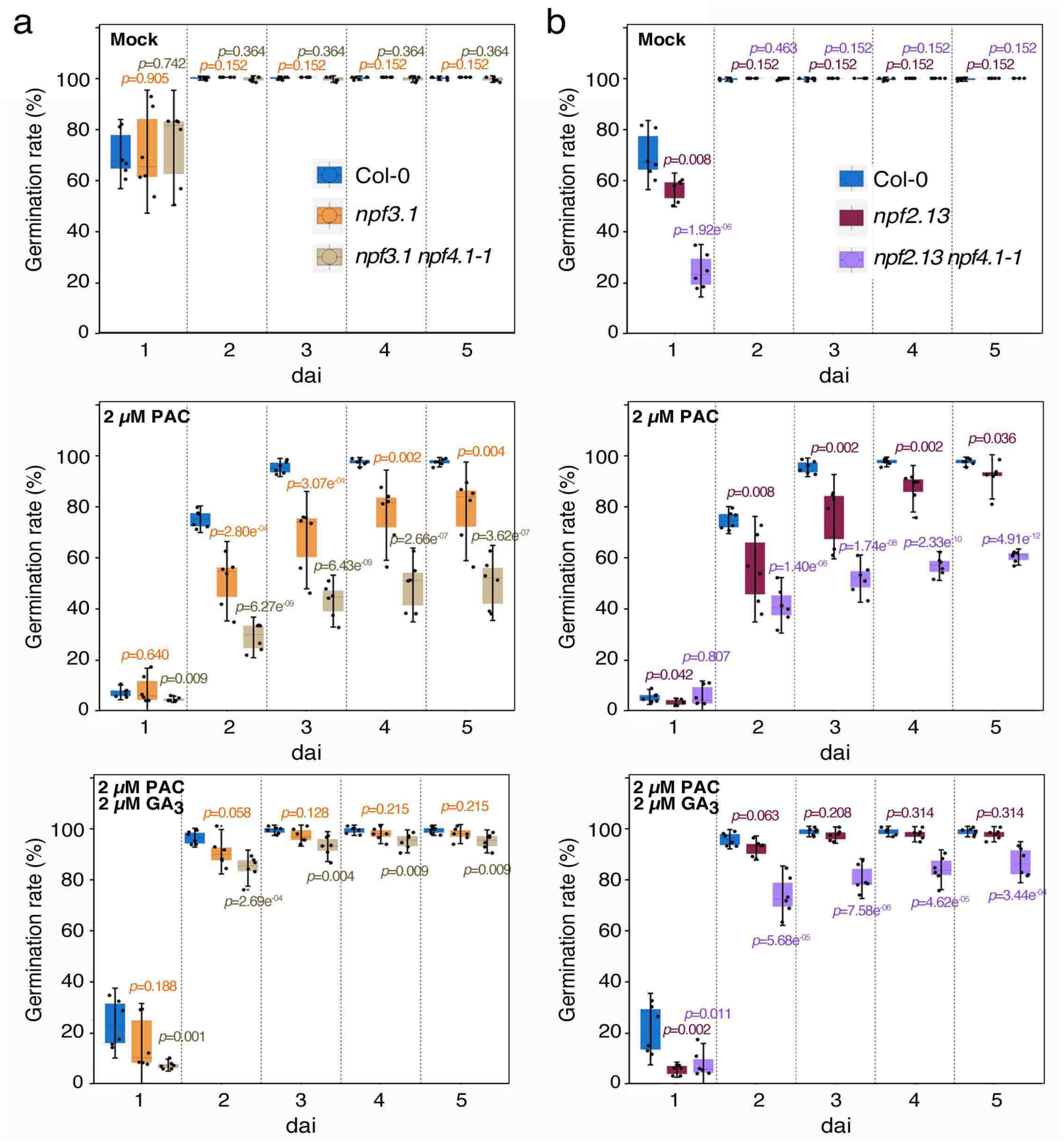
NPF3.1 and NPF2.13 regulate seed germination. (a,b) Germination of wild-type (Col-0), *npf3.1*, *npf3.1 npf4.1-1*, *npf2.13* and *npf2.13 npf4.1-1* seeds in the absence (Mock) or presence of 2 µM PAC or 2 µM PAC with 2 µM GA_3_, as indicated. Germination was determined as radicle protrusion from day 1 to day 5 after imbibition. The values are means ±SD of three seed batches of 50 to 100 seeds sown in duplicate. *P*-values are from Student’s t test for the mutants *versus* Col-0. dai, days after imbibition.

We then analyzed whether *NPF2.13* and/or *NPF3.1* have redundant activities with *NPF4.1*. To this end, we generated the *npf2.13 npf4.1-1* and *npf3.1 npf4.1-1* double mutants and tested their germination capacity. Both double mutants were more sensitive to PAC compared with *npf2.13* and *npf3.1* single mutants (Figure 7a,b), but they were as sensitive as the *npf4.1-1* single mutant (Figure 6a). Thus NPF4.1 plays the predominant role in regulating seed germination, with only small contribution from NPF2.13 and NPF3.1.

## Discussion

It has been known for decades that during germination of cereal grains, bioactive GAs produced by the embryo are released into the aleurone layers, to induce the production and secretion of α-amylase and other hydrolytic enzymes that degrade starch and various stored reserve polymers (such as carbohydrates) in the endosperm, thereby providing nutrients to the developing seedling (41). This statement is based essentially on the fact that the embryo is needed to induce starch hydrolysis in the endosperm. In barley, it has been shown that excision of the embryo prevents the accumulation of bioactive GAs in the endosperm and the activation of α-amylase (24). In addition, exogenous application of GAs substitutes the action of the embryo and induces the expression of α-amylase (23). All these observations indicate that GAs produced in the embryo induces the synthesis of α-amylase in the endosperm. However, the nature of the signal (GA mobile form), the kinetics and the transport mechanism were still unknown.

To answer these questions, we studied the transport of GAs in germinating seeds in *Arabidopsis*. Although the structure of *Arabidopsis* seeds is considerably different from that of cereals (which have a large endosperm and one or several aleurone layers), they nevertheless contain an endosperm composed of a single aleurone-type cell layer (42). And in a similar way to cereals, GAs are essential for inducing the expression of hydrolytic enzymes (cell wall remodeling enzymes, CWREs) in the endosperm (micropylar region), involved in cell wall loosening, a prerequisite for the protrusion of the radicle and therefore for the accomplishment of the germination (17).

To assess the importance of GAs for the rupture of seed coats in *Arabidopsis*, we carried out germination tests with GA-deficient seeds. As previously reported, seeds of GA biosynthesis mutants (*ga20ox1 gaox2 ga20ox3* and *ga3ox1 ga3ox2* mutants in our study) cannot germinate without exogenous GA application, unless the endosperm and testa are removed (Figure 1; 5). Thus GA-induced endosperm rupture is a key germination mechanism, although we must not neglect the important role played by GAs in radicle elongation, which provides a mechanical force that also contributes to seed coat breakdown (17, 43).

In a second step, we studied where GAs are produced in *Arabidopsis* seeds during germination. Histochemical staining with GUS reporter lines revealed that *GA3ox*, which catalyze the conversion of inactive GA_9_ into bioactive GA_4_, are essentially expressed in embryo (hypocotyl and radicle) before endosperm rupture (Figure 3). In line with the germination tests, *GA3ox1* and *GA3ox2* are the main players in the synthesis of bioactive GAs in germinating seeds (44). *GA3ox3* and *GA3ox4* are not or barely expressed in the seed. Concerning the *GA20ox* (which synthetize GA_9_), GUS activity staining showed that *GA20ox1* and *GA20ox2* are expressed exclusively in the embryo, while *GA20ox3* is expressed in both embryo and endosperm (Figure 3). Thus, our results indicate that while GA biosynthetic intermediates are produced in embryo and endosperm, the active GAs are predominantly synthetized in embryo, in agreement with previous studies in cereals (24).

If the embryo is the main source of bioactive GA production in germinating seeds, these GAs must necessarily be transported to the endosperm to induce the expression of *CWRE* genes. To indirectly visualize the transport of GAs from the embryo to the endosperm, we undertook a strategy similar to that used in cereals, which consisted of monitoring the induction of α-amylase production in aleurone cells (23). Here in *Arabidopsis* seeds, we monitored in a series of experiments the spatio-temporal expression pattern of *EXPA2*, a GA-induced endosperm-specific CWRE marker (17). Through an “embryo-bedding assay” combined with the application of exogenous GA_4_, we first confirmed that GA_4_ is indeed transported from the embryo to the endosperm in *Arabidopsis* seeds (Figure 4). However, we cannot exclude the possibility that other forms of GAs are also transported, as it is the case for GA_9_ in flowers and GA_12_ over long distances (45, 46). Nevertheless, given that *GA3ox* are barely expressed in the endosperm, the eventual transport of GA intermediates would have little impact on *CRWE* induction. This was confirmed in our study, where application of GA_9_ to dissected seed coats did not induce the expression of *EXPA2* in endosperm (Figure 4c). We then showed that embryo-derived GA_4_ begins to move in the endosperm shortly before 9 hours of imbibition, which is consistent with the expression level of *GA3ox1* and *GA3ox2* in the embryo, at this stage.

To broaden our knowledge on the embryo-to-endosperm transport of GA_4_, we then searched for the protein carriers. Although GAs are weak acids that can in theory diffuse and enter into the cells freely, several recent studies have identified transporters that orientate the flow of GAs from production to action sites (Binenbaum et al., 2018). For example, this is the case for NPF3.1 that concentrates GAs in the elongating endodermal cells of the root (25). In our study, we showed that NPF4.1, which displays high GA import activities (25, 39), is strictly expressed in the endosperm from the first few hours after seed imbibition, thus, concomitant with the synthesis of GA_4_ in embryo. Consistent with the premise that NPF4.1 facilitates the transport of GA_4_ from the embryo to the endosperm, we showed that induction of *EXPA2* is restrained in *npf4.1* imbibed seeds (Figure 6c). In addition to NPF4.1, we identified two other GA transporters in germinating seeds, NPF2.13 and NPF3.1. These two transporters are mostly expressed in cotyledons and radicle, respectively, suggesting a role in GA-mediated embryo growth (Figure 5b). Noteworthy, previous studies have revealed that NPF4.1, NPF2.13 and NPF3.1 have dual GA and ABA transport activities (25, 38, 39, 47). Since these two hormones play antagonistic role in germination (1), the regulation of their activity towards GA and ABA transport must be finely controlled. This is at least the case for NPF4.1, for which ABA transport activity is inhibited by GAs (47).

To further substantiate the biological function of these three transporters, we finally analyzed the germination capacity of each respective mutant. Germination tests were carried out with stratified non-dormant seeds to attenuate the effects of ABA. These assays showed that *npf4.1*, *npf3.1* and *npf2.13* mutant seeds are hypersensitive to PAC, and that addition of GAs suppresses the effect of PAC, hence confirming their role in GA-mediated seed germination (Figure 6 and 7). Meanwhile, the near absence of germination defects in *npf4.1*, *npf3.1* and *npf2.13* mutants under control conditions strongly suggests the existence of functionally redundant GA transporters in seeds. A previous study reported that mutation of two GA transporters, SWEET13 and SWEET14, previously linked to sugar transport, increased seed resistance to PAC (30). In our growth conditions, we couldn’t observe a similar trend with the *sweet13 sweet14* double mutant; different seed age, PAC concentration or germination evaluation method could explain this discrepancy. So far, no other GA transporters have been shown to regulate GA-mediated physiological processes in seeds.

In summary, based on our results, we can build the following model (Supplementary Figure 7). A few hours after seed imbibition, GA3ox1 and GA3ox2 rapidly produce bioactive GA_4_ in the hypocotyl. NPF3.1 and NPF2.13, which accumulate very early in the radicle and cotyledons, likely contribute in the distribution of GA_4_ in these compartments to induce cell expansion and thereby the growth potential of the embryo. Concomitantly, NPF4.1 facilitates the transport of embryo-derived GA_4_ into the endosperm, hence inducing the expression of *CWREs* (including *EXPA2*) involved in the rupture of the seed coats, a prerequisite for the protrusion of the radicle.

## Acknowledgements

We thank Tp. Sun for providing seeds of *ga3ox1 ga3ox2* and *pGA3ox:GUS* transgenic lines; P. Hedden for *ga20ox1 ga20ox2 ga20ox3* and *pGA20ox:GUS* lines; L. Onate-Sanchez for *pEXPA2:GUS* and S. Ferrario-Mery for *npf3.1* (Salk_130095). We thank the personnel of Institut de Biologie Moléculaire des Plantes (IBMP) facilities (UPR2357, Strasbourg) for assistance with microscopy, gene expression analysis and plant care. We thank E. Lechner, S. Ferrario-Méry and colleagues from Gibberellins and plant adaptation team (IBMP) for fruitful discussions on this work. This work was supported by the Centre National de la Recherche Scientifique and the French ministry of research and higher education.

## Author contributions

All experimental work, except bioinformatics analysis, was conducted by M.S.J., L.S.A. and P.A.; bioinformatics (Supplementary Table 1) was performed by D.P.; M.S.J., J.M.D. and P.A. designed the experiments; J.M.D. and P.A. supervised the work; M.S.J., J.M.D. and P.A. wrote the paper.

## Methods

### Plant material

Mutants and transgenic lines were in the Columbia-0 (Col-0) background. *NPF4.1-1* (WiscDsLox434A10) and *NPF4.1-2* (SAIL_832_E12.C) were supplied by the Nottingham *Arabidopsis* Stock Centre. *npf3.1* (SALK_130095), *npf2.13* (SALK_022429), *ga20ox1 ga20ox2 ga20ox3*, *ga3ox1 ga3ox2*, *pNPF2.13:GUS*, *pGA20ox:GUS*, *pGA3ox:GUS* and *pEXPA2:GUS* were already described (17, 29, 31, 32, 38, 44). PCR genotyping for selecting homozygous lines was performed using specific primers listed in Supplementary Table 2. To generate *npf4.1-1 npf2.13* and *npf4.1-1 npf3.1* double mutants, respective single mutants were crossed, and F3 homozygous plants were selected by PCR genotyping. To generate *pNPF4.1:GUS* and *pNPF3.1:GUS*, the promoter of *NPF4.1* (1,5-kb fragment) and *NPF3.1* (2-kb fragment) were PCR amplified from Col-0 genomic DNA with specific primers listed in Supplementary Table 2, inserted into pDNOR221 (Invitrogen) and then recombined into pGWB633 (48). To generate *p35S:NPF4.1-GFP* and *p35S:NPF4.1-HA*, coding sequence of *NPF4.1* was PCR amplified using specific primers listed in Supplementary Table 2, inserted into pDONR/ZEO (Invitrogen) and then recombined into pB7FWG2 and pGWB14, respectively (49, 50). The different constructions were introduced into *Agrobacterium* GV3101 and then transformed into *Arabidopsis* Col-0 by the floral dip method.

### Growth conditions

Given that the timing of seed germination is modulated by different environmental and internal factors including the age and dormancy level of the seeds, all experiments in this study were performed with a standard procedure. To attenuate these effects and obtain reproducible and comparable results, sterilized seeds (aged between 6 and 12 months) were stratified at 4°C for 48 hours on plates containing half Murashige-Skoog (½ MS) medium (Duchefa Biochemical), 0.5% sucrose and 0.8% agar (Sigma-Aldrich). After stratification, plates were transferred in growth chamber under long day photoperiod (16h light at 22°C / 8h dark at 22°C, irradiance conditions of about 70 µmol m^-2^ s^-1^). To germinate, GA-deficient seeds (*ga20ox1 ga20ox2 ga20ox3* and *ga3ox1 ga3ox2* mutants) were pre-treated during stratification with 10 µM GA_3_ (Olchemim), unless otherwise specified. For chemical application, GAs (Olchemim) and PAC (Sigma-Aldrich) were directly added to the ½ MS agar medium at concentrations indicated in the figure legends.

### Germination tests

For the germination assays, non-dormant mature seeds were disposed on ½ MS agar plates containing appropriate concentration of GA_3_ and PAC as indicated in the figure legends. Germination, scored every day, was defined as radicle protrusion, except in Figure 1b, where it was defined as cotyledon opening 4 days after imbibition.

### Embryo bedding assay

The assay was carried out in the same way as the “seed coat bedding” assay previously described (37). Briefly, non-dormant Col-0, *ga3ox1 ga3ox2* and *pEXPA2:GUS* seeds were dissected one hour after imbibition starting point. Separated *pEXPA2:GUS* seed coats were disposed on a layer of Col-0 or *ga3ox1 ga3ox2* embryos disposed on a 70 µm cell strainer (Falcon) directly placed on ½ MS agar plate. 24 hours after incubation, EXPA2:GUS staining was conducted on the seed coats.

### GUS analysis

GUS staining was performed on dissected embryos and seed coats. Samples were fixed in 90% acetone for 15 min in ice, infiltrated in a GUS solution composed of 200 µg/ml (for embryos) or 400 µg/ml (for seed coats) 5-bromo-4-chloro-3-indolyl-ß-D-glucoronide (X-GLUC), 50mM sodium phosphate pH 7.0, 1 mM potassium ferricyanide, 1mM potassium ferrocyanide, 10 mM ethylenediamine tetraacetic acid (EDTA), 0.01% triton X-100 for 15 min, and incubated at 37°C overnight. The reaction was stopped with 70% ethanol. The samples were fixed in a solution of 100% ethanol and glacial acetic acid (3:1) for a day, and then washed twice, in water. Seed coats were destained in a 2.5% sodium hypochlorite solution until whitening.

### Gene expression analysis

Total RNAs from whole seeds, separated embryos or seed coats, were extracted with the RNeasy Plus Mini kit (Qiagen), according to manufacturer’s procedure. RNA samples were reverse-transcribed into cDNA with Superscript IV reverse transcriptase (Invitrogen). qRT-PCR was performed using gene-specific primers (listed in Supplementary Table 2) in a total volume of 10 µL SYBR Green Master mix (Roche), on a Lightcycler LC480 apparatus (Roche), according to manufacturer’s instructions. The PCR programme was as following: an initial DNA denaturation at 95°C for 5 min; 45 cycles including a denaturation step at 95°C for 10 sec, an annealing step at 60°C for 15 sec and an extension step at 72°C for 15 sec; and a melting curve from 55°C to 95°C. *SAND* (AT2G28390) and *EXP* (AT4G26410) genes were used as internal reference genes. *ABI4* (AT2G40220) and *EXPA2* (AT5G05290) genes were used as embryo and seed coat specific markers, respectively. Expression level of each gene was calculated using Lightcycler 480 software, release 1.5.0 SP3.

### Observation of GFP fluorescence

NPF4.1-GFP fluorescence from root of 7-day-old *p35S:NPF4.1*-GFP seedlings was observed with a Zeiss LSM780 inverted confocal laser microscope with 40x objectives. Root cell walls were stained with propidium iodide (10 µg/ml, Sigma-Aldrich).

### Bioinformatics

RNA-seq data reported in Supplementary Table 1 (extracted from dataset GSE224926; 34) was processed using the nf-core RNA-seq pipeline version 3.10 (51) using the default options (https://nf-co.re/rnaseq/3.10). Read alignment was performed on TAIR10 genome with STAR and reads quantification was done with Salmon. Then a differential expression analysis was done in R using the DESeq2 package (52).

### Statistical analyses

Statistical analyses were performed using RStudio package v.1.2.1335 (www.rstudio.com). The different analyses (Student test, 1 way-Anova and Tukey comparison) were made using a significance threshold of 5% (p < 0.05).

---

**Supplementary Figure 1.**
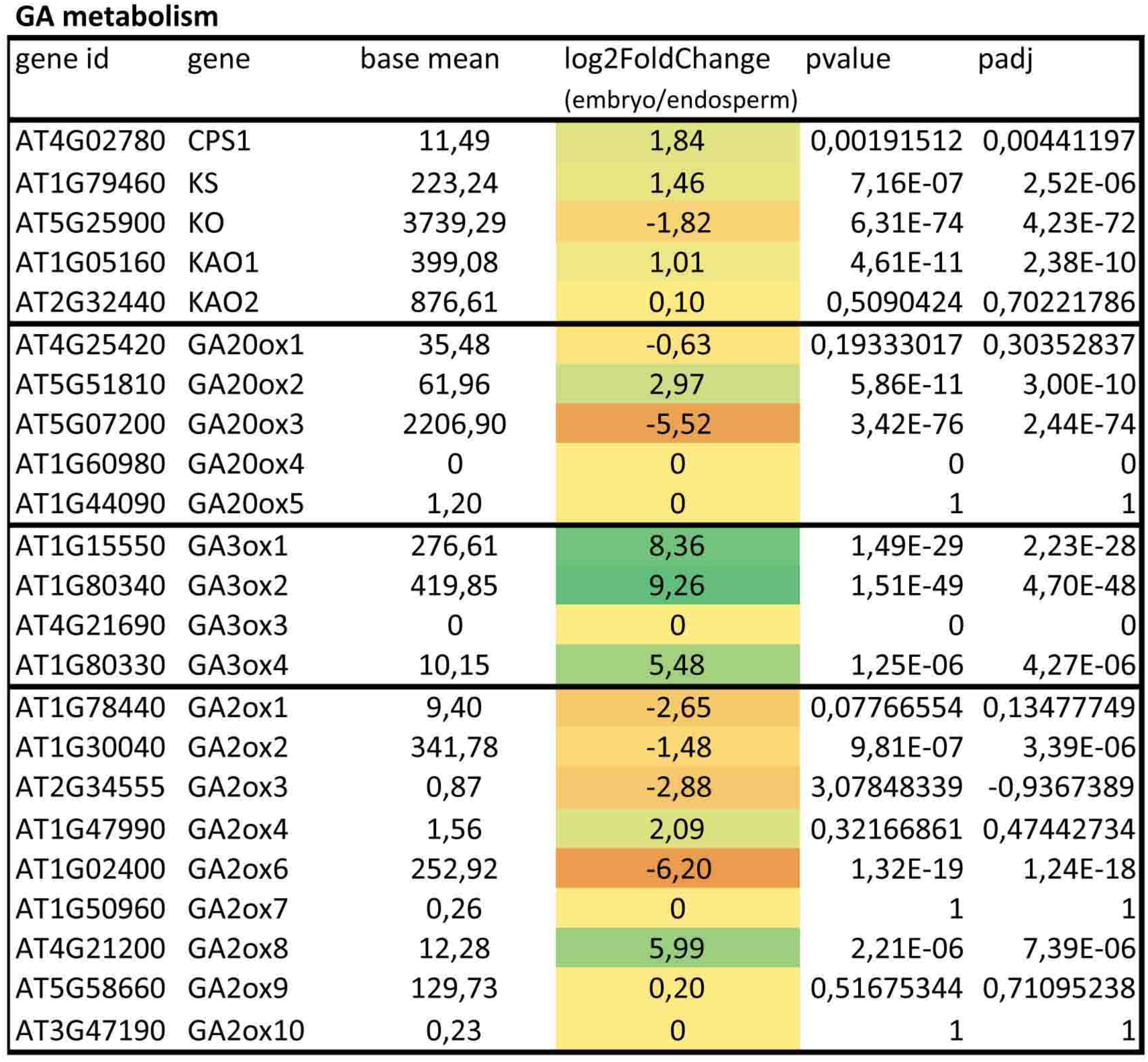
Transcriptional behavior of GA metabolism genes in *Arabidopsis* embryo and endosperm cultured for 24 hours. Heatmap representation of the log2 fold-change (embryo/endosperm) of GA metabolism genes. Green (induction) and red (repression) color intensity is proportional to log2 fold-change value. Base mean expression, pvalue and padj of each GA metabolism gene are indicated. Complete gene expression analysis is provided in Supplementary Table 1. Data were extracted and reanalyzed from a previous RNA-seq analysis performed by Piskurewicz et al. 2023 (34) (dataset GSE224926), carried out on dissected wild-type embryos and endosperms incubated for 24 hours at 30°C. For further details, see Methods.

**Supplementary Figure 2.**
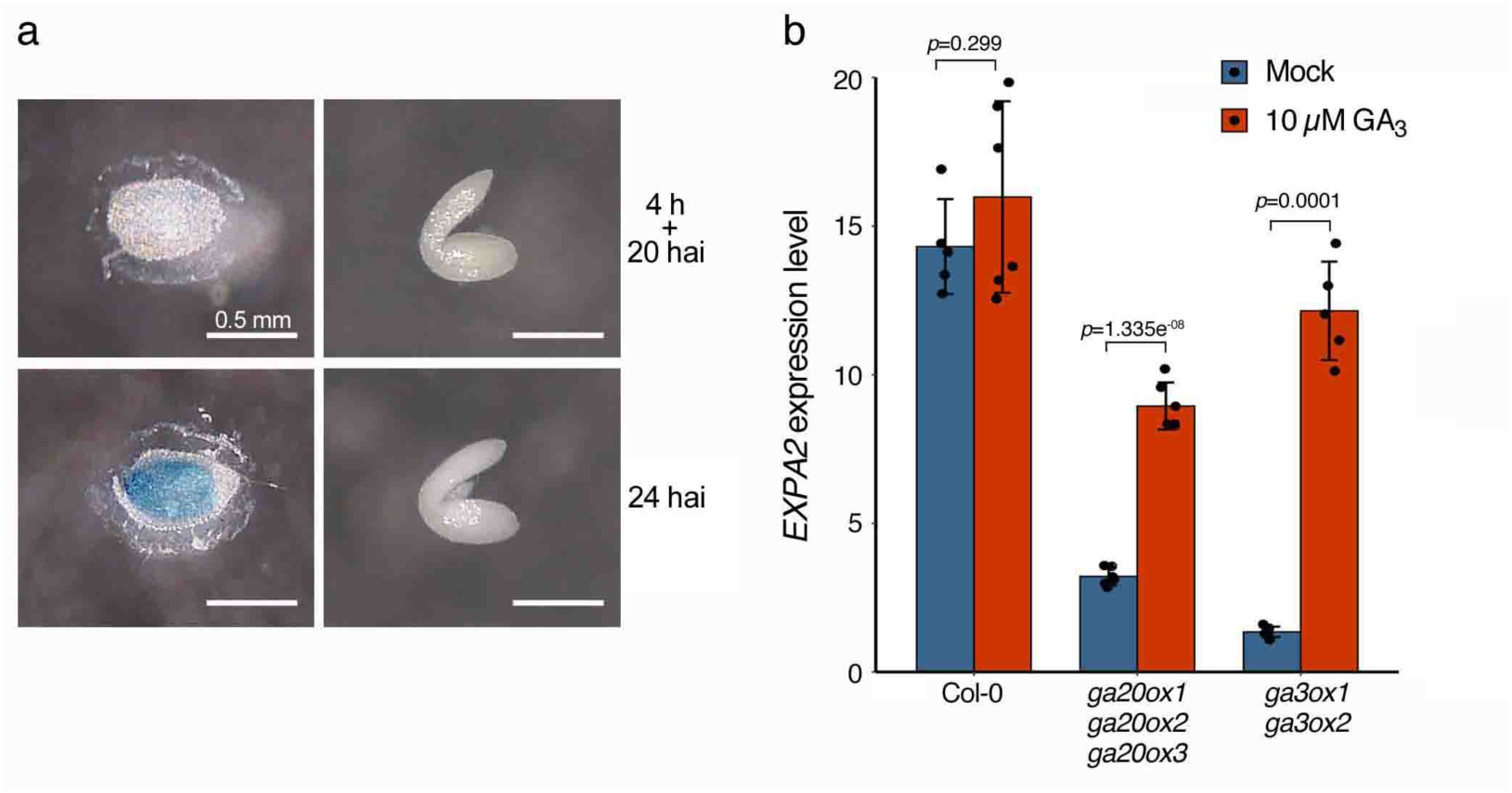
*EXPA2* is a GA-inducible gene expressed in endosperm. (a) EXPA2:GUS activity in dissected embryo and seed coats of 24 hours imbibed *pEXPA2:GUS* seeds, isolated 4 hours or 24 hours after imbibition. (b) Expression level of *EXPA2* in wild-type (Col-0), *ga20ox1 ga20ox2 ga20ox3* and *ga3ox1 ga3ox2* mutant seeds imbibed for 24 hours with 10 µM GA_3_ and untreated controls (Mock). Data are means ±SD of two independent replicates with two to three technical replicates. *P*-values are from Student’s t test.

**Supplementary Figure 3.**
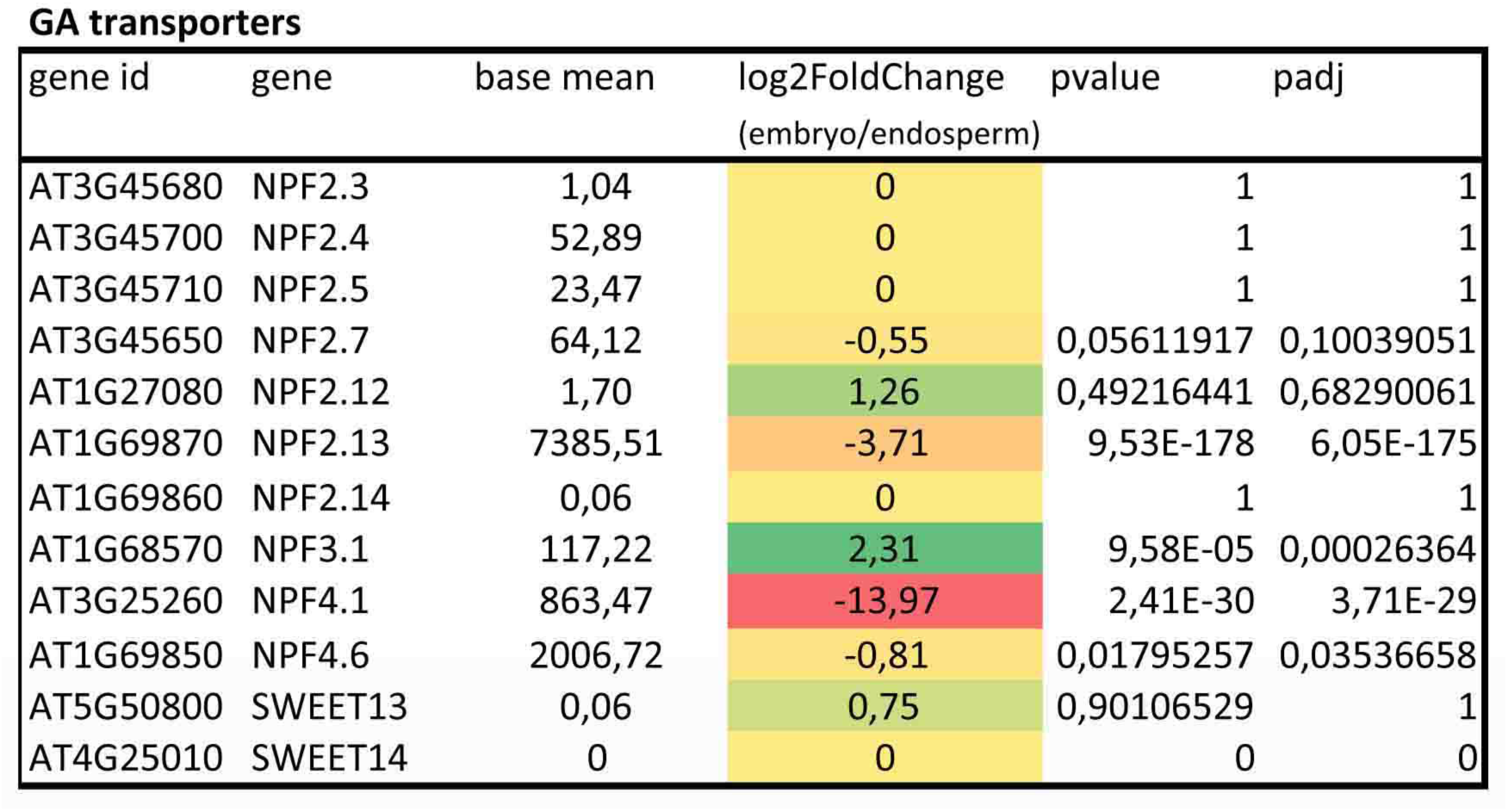
Transcriptional behavior of GA transporter genes in dissected embryo and endosperm cultured for 24 hours. Heatmap representation of the log2 fold-change (embryo/endosperm) of GA transporter genes. Green (induction) and red (repression) color intensity is proportional to log2 fold-change value. Base mean expression, pvalue and padj of each GA transporter gene are indicated. Complete gene expression analysis is provided in Supplementary Table 1. Data were extracted and reanalyzed similarly as in Supplementary Figure 1, from a previous RNA-seq analysis performed by Piskurewicz et al. 2023 (34) (dataset GSE224926), carried out on dissected wild-type embryos and endosperms incubated for 24 hours at 30°C.

**Supplementary Figure 4.**
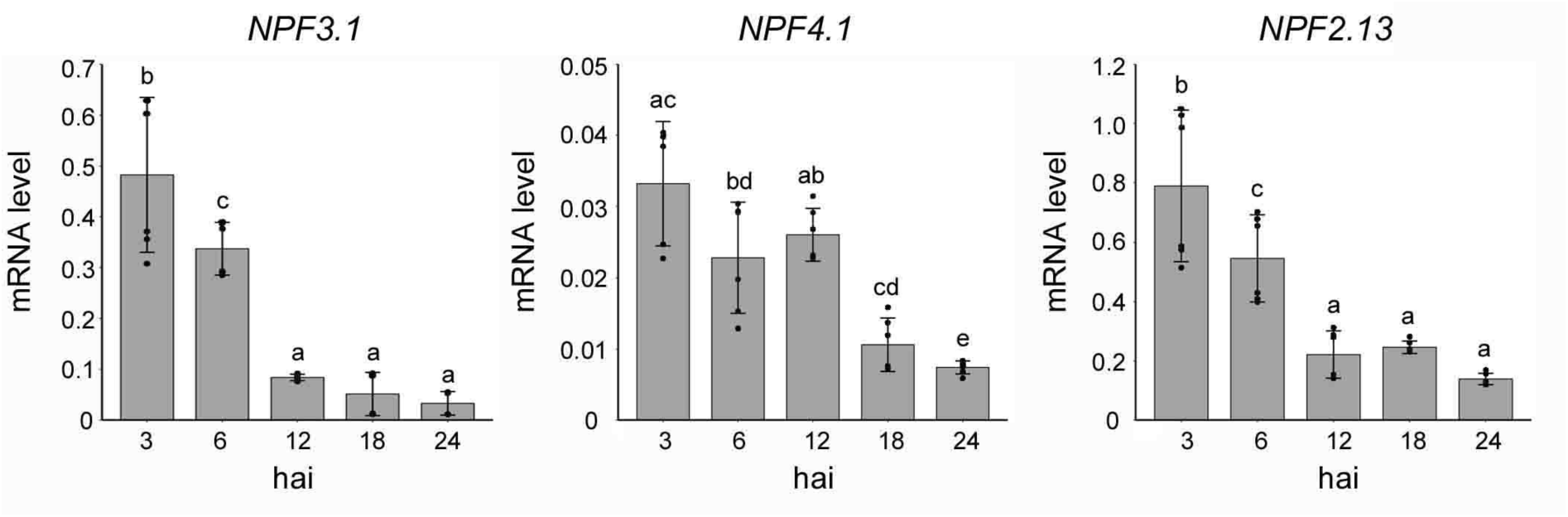
Temporal expression of *NPF3.1*, *NPF4.1* and *NPF2.13* during seed germination. Expression level of *NPF3.1*, *NPF4.1* and *NPF2.13* in imbibed seeds for 3, 6, 12, 18 and 24 hours after imbibition. The values are means ±SD of two independent replicates with two to three technical replicates. Different letters denote significant differences (p<0.05) using one-way ANOVA with Tukey’s test for multiple comparisons.

**Supplementary Figure 5.**
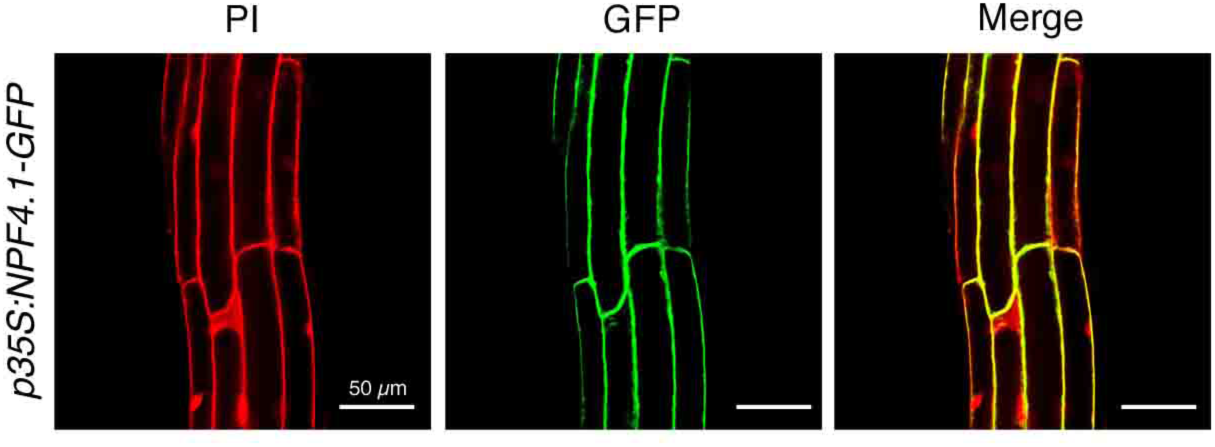
Subcellular localization of NPF4.1. Confocal imaging of 7-day-old root epidermal cells expressing *35S:NPF4.1-GFP*. Cell wall was stained with propidium iodide (PI). The experiment was repeated twice with similar results. Scale bars, 50 µm.

**Supplementary Figure 6.**
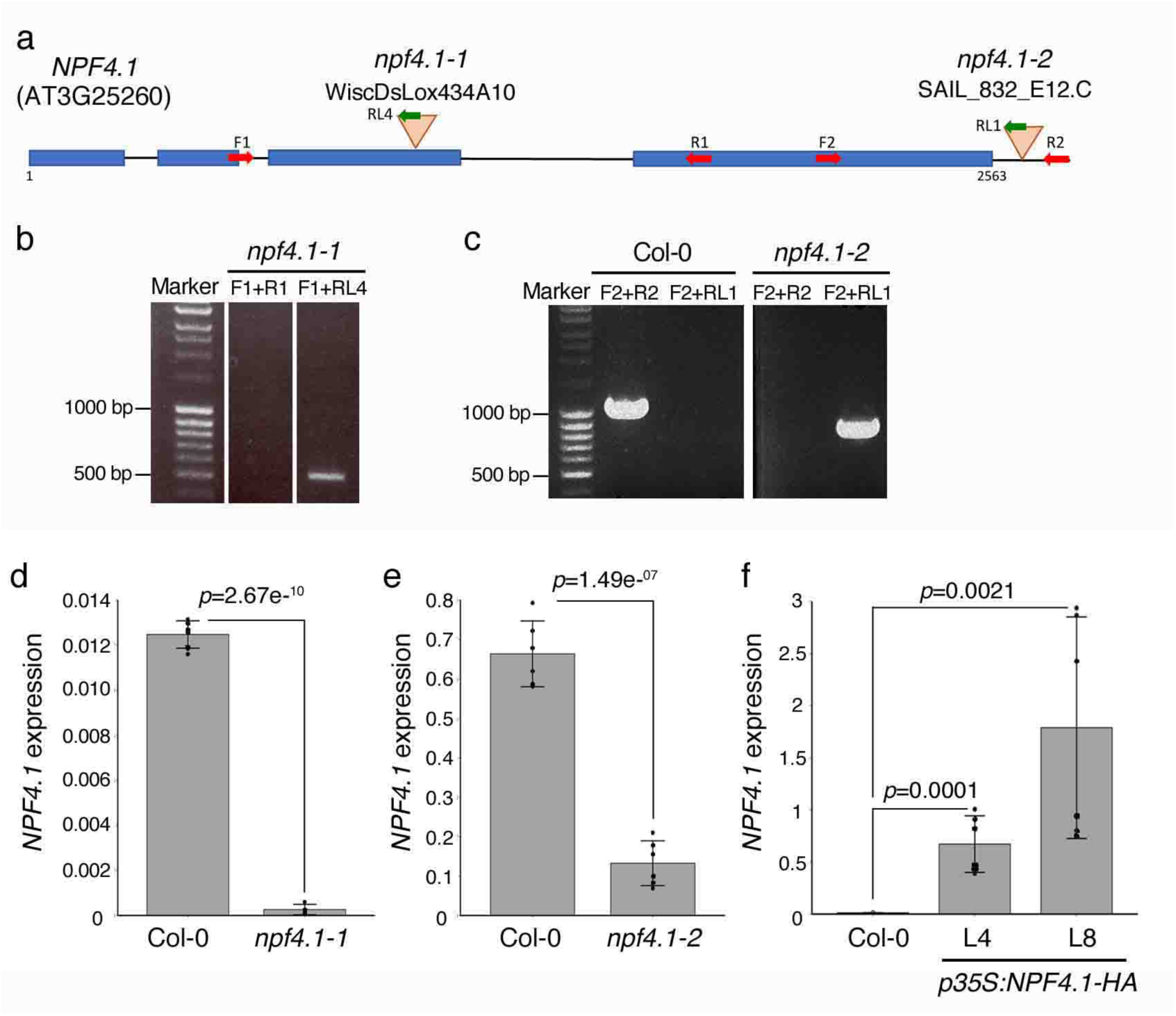
Characterization of *npf4.1* mutants and *p35S:NPF4.1-HA* transgenic lines. (a) Gene model of *NPF4.1* showing the T-DNA insertion site of mutant alleles. Horizontal boxes denote exons and horizontal lines denote introns. Red and green arrows indicate the position of oligonucleotides used for genotyping. (b,c) PCR genotyping confirming that *npf4.1-1* (b) and *npf4.1-2* (c) are two homozygous mutants. Marker, DNA ladder. The experiment was repeated twice with similar results. (d,e) RT-qPCR indicating that *npf4.1-1* is a null mutant allele while *npf4.1-2* is a knock-down mutant allele. (d) *NPF4.1* expression in 24 hours imbibed wild-type and *npf4.1-1* seeds. Data are means ±SD of two experiments with three technical replicates. *P*-value is from Student’s t test. (e) *NPF4.1* expression in 24 hours imbibed wild-type and *npf4.1-2* seeds. Data are means ±SD of two experiments with three technical replicates. *P*-value is from Student’s t test. (f) *NPF4.1* expression in 24 hours imbibed wild-type and *p35S:NPF4.1-HA* line 4 (L4) and line 8 (L8) seeds. Data are means ±SD of two experiments with three technical replicates. *P*-values are from Student’s t test.

**Supplementary Figure 7.**
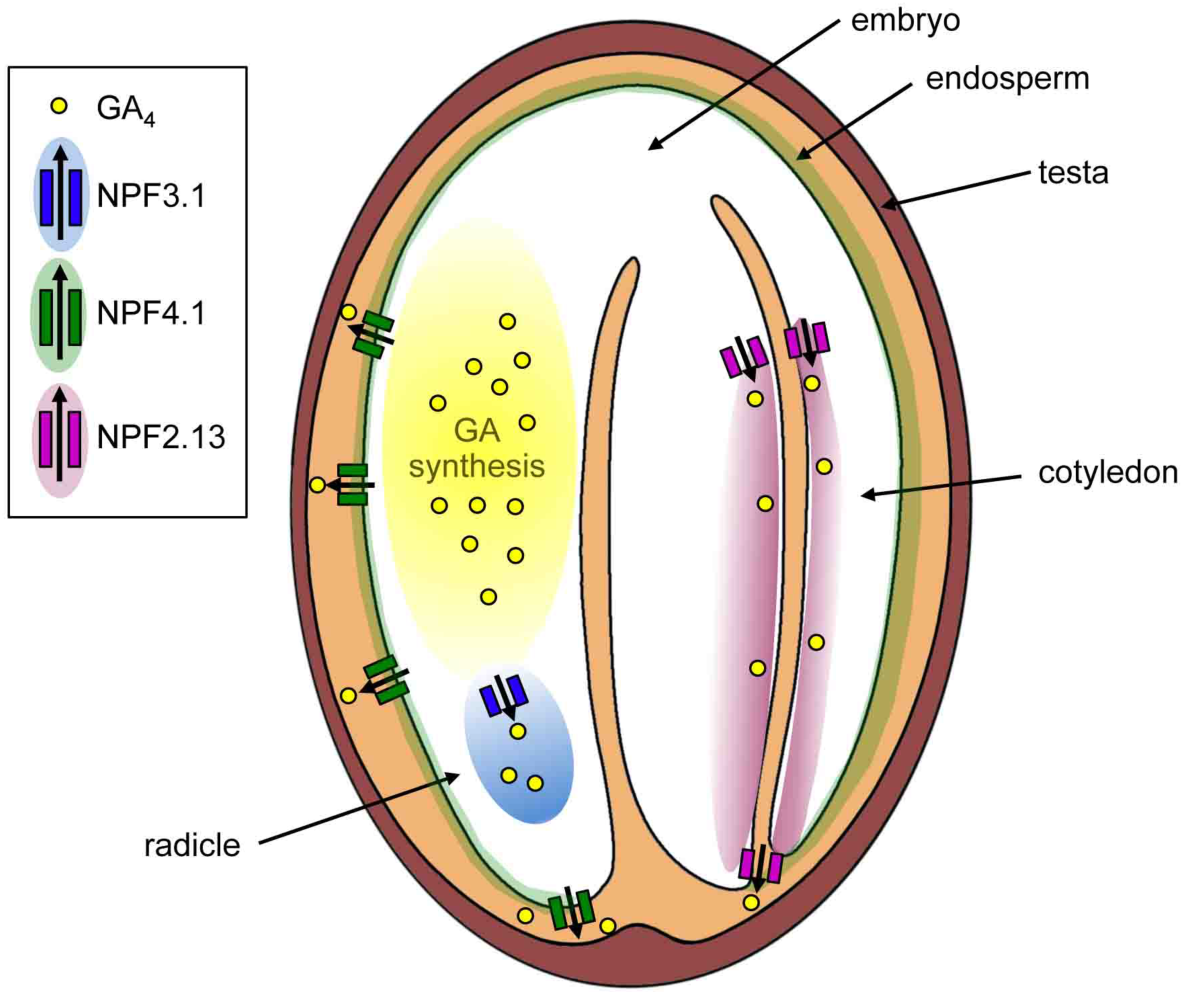
Proposed model illustrating the function of NPF4.1, NPF3.1 and NPF2.13 during seed germination. In the first few hours after imbibition, GA3ox1 and GA3ox2 synthetize GA_4_ in the hypocotyl embryo. NPF3.1 and NPF2.13, which are respectively localized in radicle and cotyledons, orientate the movement of GA_4_ in these organs to induce cell expansion, increasing the overall growth potential of the embryo. Concomitantly, NPF4.1, which is localized in the plasma membrane of endosperm cells, facilitates the transport of embryo-derived GA_4_ to the endosperm to induce the expression of *CWREs* (including *EXPA2*) that activate cell expansion and separation to allow radicle protrusion.

**Supplementary Table 1. Transcriptome of dissected wild-type embryos and endosperms incubated for 24 hours at 30°C**

Data were extracted and reanalyzed from a previous RNA-seq analysis performed by Piskurewicz et al. 2023 (34) (dataset GSE224926). For more information, see Methods.

**Supplementary Table 2. List of the oligonucleotides used in the study**

